# CNpare: matching DNA copy number profiles

**DOI:** 10.1101/2021.09.28.462193

**Authors:** Blas Chaves-Urbano, Bárbara Hernando, Maria J Garcia, Geoff Macintyre

## Abstract

Selecting the optimal cancer cell line for an experiment can be challenging given the diversity of lines available. Cell lines are often chosen based on their tissue of origin, however, the results of large-scale pan-cancer studies suggest that matching lines based on molecular features may be more appropriate. Existing approaches are available for matching lines based on gene expression, DNA methylation or low resolution DNA copy number features. However, a specific tool for computing similarity based on high resolution genome-wide copy number profiles is lacking. Here, we present CNpare, which identifies similar cell line models based on genome-wide DNA copy number. CNpare compares copy number profiles using four different similarity metrics, quantifies the extent of genome differences between pairs, and facilitates comparison based on copy number signatures. CNpare incorporates a precomputed database of 1,170 human cancer cell line profiles for comparison. In an analysis of separate cultures of 304 cell line pairs, CNpare identified the matched lines in all cases. CNpare provides a powerful solution to the problem of selecting the best cell line models for cancer research, especially in the context of studying chromosomal instability.

## Introduction

Immortalised cancer cell lines are an integral part of cancer research and the development of new therapies^1,2^. Cell lines are often selected based on their tissue of origin. However, a suite of approaches are emerging which facilitate appropriate cell line selection based on molecular features such as gene expression, DNA methylation, and genomics^3–13^. A subset of these approaches perform DNA copy number based comparison at different resolutions including gene-level copy number, chromosome arm copy number, ploidy, or genome doubling status^9^. However, a specific tool for computing similarity based on genome-wide copy number is lacking.

Sophisticated tools using evolutionary models can be used to compare genome-wide copy number between tumours from the same patient^14^ and cells within a tumour^15^. However, tumours from different patients do not share an evolutionary relationship, therefore more traditional similarity metrics can be used. Here, we present CNpare, which can be used to: 1) calculate similarities between genome-wide copy-number profiles; 2) quantify the extent of genome differences between copy-number profiles; and 3) compare profiles based on copy-number signatures^16^ (Figure 1).

**Figure 1.**
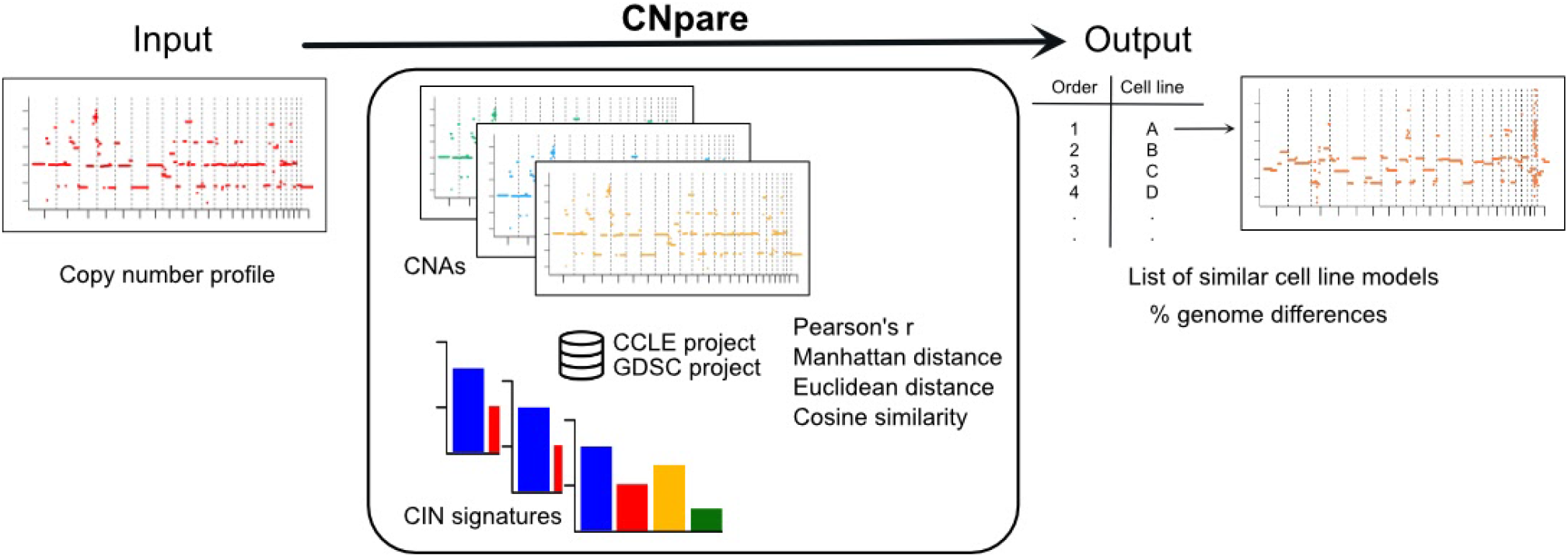
Overview of the CNpare method. This schematic provides a high-level overview of CNpare’s workflow and computation. The user inputs an absolute copy number profile (left) and CNpare compares this to a precomputed database of cell line copy number from the CCLE and GDSC projects, using a series of different comparison metrics (centre). Output is in the form of a list of cell lines ranked based on the strength of match to the input profile. Included in the output is a graphical representation of differences between the genomes and an estimate on the percent genome difference.

## Results

### Assessing performance

To assess the performance of CNpare, we used separate cultures of the same cell lines profiled as part of the CCLE and GDSC projects (304 pairs). We observed the ability of CNpare to identify, for each GDSC line, the correctly matched line in the CCLE database (Figure 2). In 100% of cases the top hit for each GDSC line was the matched line in the CCLE database. 87% (265 cells) were matched correctly across all similarity metrics, with the remaining 13% being matched correctly only by Pearson correlation and cosine similarity (not Manhattan and Euclidean distance). For these cases, the ploidy estimated by ASCAT differed between the CCLE and GDSC cultures causing comparison based on Manhattan and Euclidean distances to return the wrong cell line. As Pearson correlation and Cosine similarity are normalized measures, they successfully identified the correct profile independent of the difference in ploidy status between the cultures. This difference in metric performance demonstrates the choice of metric can determine whether the comparison is ploidy aware or agnostic. As such, we developed a guide on which metrics to use for different circumstances (Table 1).

**Table 1.** Comparison metrics and recommended use.

| Similarity metrics | Properties | Recommendations |
| --- | --- | --- |
| Pearson's r | Normalized-based metric<br>• Invariant to scaling, thus magnitude is ignored<br>• Invariant to location shifts of data values | Identification of similar profiles regardless of their ploidy status. Preferable to cosine similarity for the case of highly fragmented profiles |
| Cosine similarity | Normalized-based metric<br>• Invariant to scaling, thus magnitude is ignored<br>• Used for high-dimensional data | Identification of similar profiles regardless of their ploidy status |
| Manhattan distance | Magnitude-based metric<br>• Affected by the feature units<br>• Used for high-dimensional data | Identification of profiles with similar focal events and ploidy status. Preferable to euclidean distance for the case of highly fragmented profiles |
| Euclidean distance | Magnitude-based metric<br>• Affected by the feature units<br>• Used for low-dimensional data | Identification of profiles with similar focal events and ploidy status |

**Figure 2.**
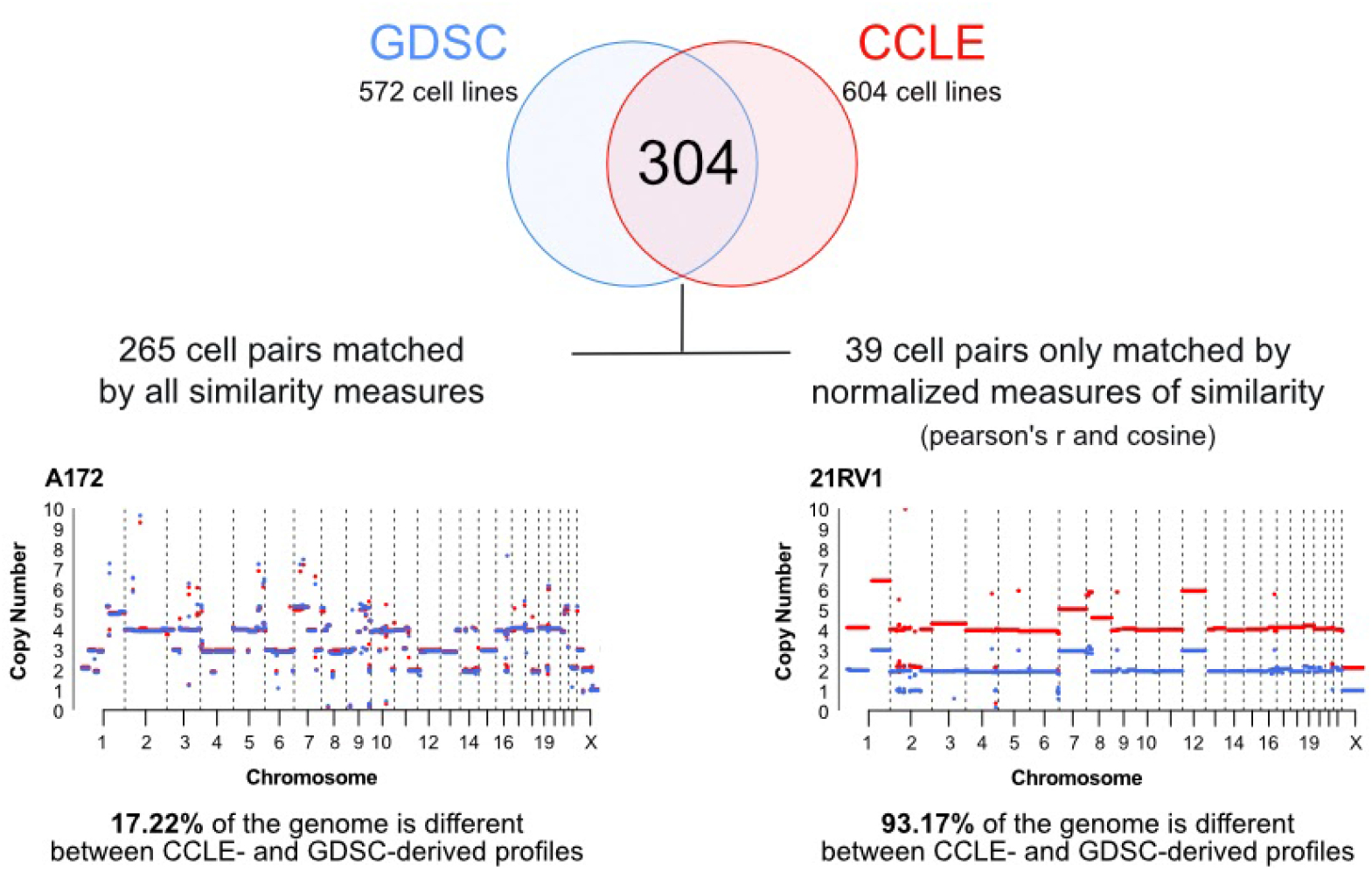
Summary of CNpare performance when used to correctly match 304 GDSC cell lines within the CCLE database. The copy number plot on the left shows an example of a GDSC culture cell line copy number profile (blue) matched with its CCLE culture profile top hit (red) across all similarity metrics. In total, 265 of the 304 lines matched across all metrics. The copy number plot on the right is an example of one of the 39 cell line pairs which did not match for Manhattan and Euclidean distance (but did match for Pearson’s r and Cosine similarity). In this case, these cell lines have different copy number ploidy fits but similar copy number changes.

### Comparison against existing approaches

CNpare is not designed to be superior to existing approaches but rather provide an alternative for choosing matched cell lines which is based on genome-wide DNA copy number. A head-to-head comparison with existing approaches is difficult due to a lack of a gold standard for matching similar cell lines. However, in an attempt to perform a comparison with existing approaches, we rederived certain cell line data that were representative of those used by existing approaches including: gene based copy number, chromosome arm based copy number, ploidy and gene expression. Using these data to compare (Pearson’s r) the cell line culture pairs analysed above, we found that gene based copy number matched 100% of the cell lines correctly, arm based copy number 81%, ploidy 82%, and gene expression 38%. This suggests that higher resolution DNA copy number based matching provides a robust method of identifying similar cell line models.

### Case study: matching an ovarian cancer cell line

Using the high-grade serous ovarian cancer cell line OVKATE, we sought the next most representative cell line model in the CNpare database.

First, we directly compared the OVKATE copy number profile with the database and the cell line which had the most similar copy-number profile was PANC0203 (Pearson’s r=0.44, Manhattan=0.97, with 52% genome difference, Figure 3). Despite this line being derived from a pancreatic adenocarcinoma, it showed highly correlated gene expression (r=0.79, p=0.02) and DNA methylation status (r=0.79, p=0.03).

**Figure 3.**
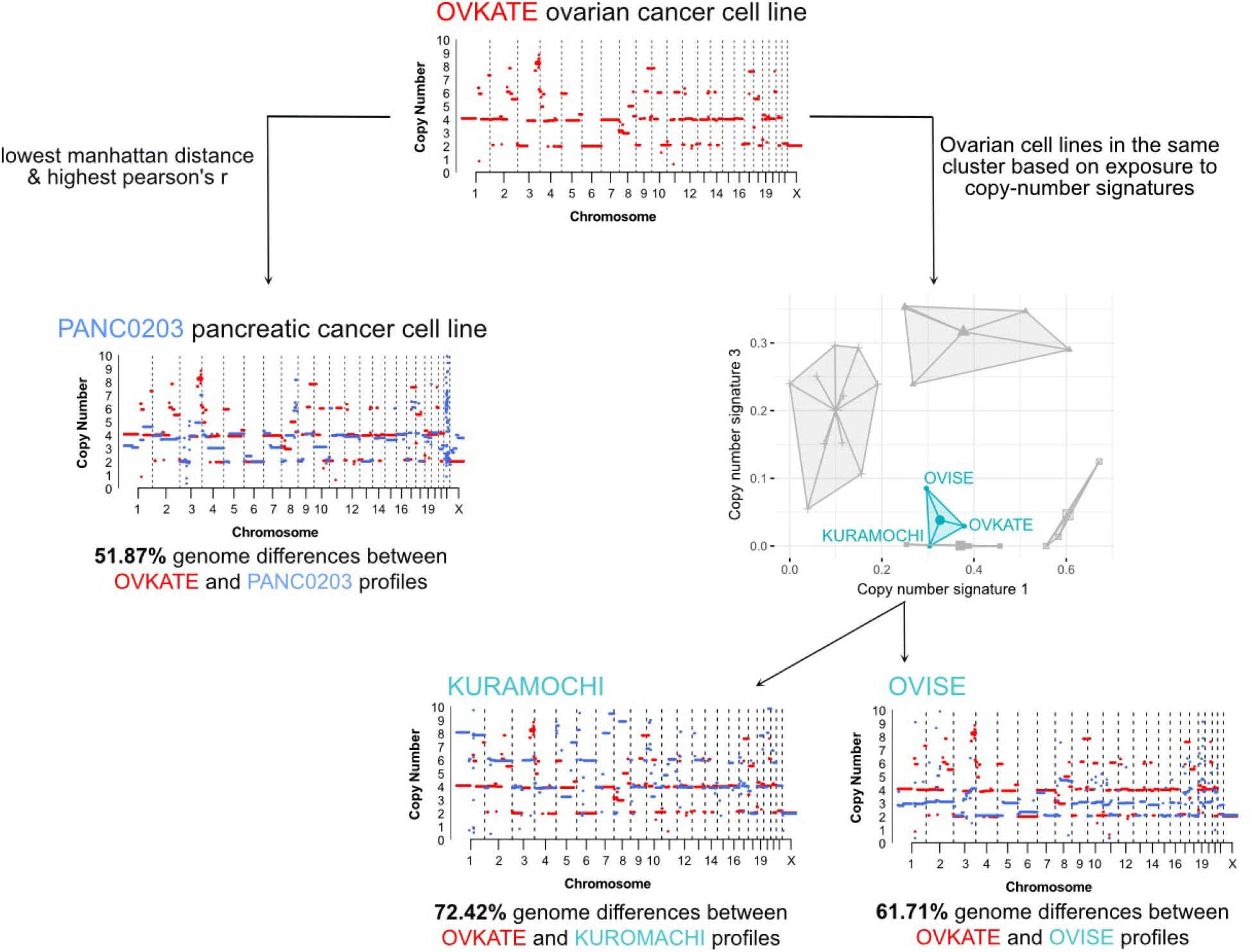
Results of matching the OVKATE cell line with it’s most representative cell line in the database using CNpare. The top plot shows the copy number profile of the OVKATE ovarian cancer cell line used as input to CNpare. On the left side, the copy number profile of the top hit found using Pearson’s r and Manhattan distance is displayed in blue, along with the OVKATE line in red. Underneath the percentage genome difference between the profiles is listed. On the right hand side, the results of matching the OVKATE cell line based on copy number signatures is displayed. The top plot shows the results of clustering all ovarian cancer cell lines based on 7 copy number signatures. For visualisation purposes, the two signatures with the highest variation across cluster means are shown. Each large dot represents the cluster centroid and each small dot represents a cell line. The cluster containing the OVKATE cell line is colour coded in turquoise. The copy number plots below show the differences in copy number profiles between OVKATE and the matching KURAMOCHI and OVISE cell lines.

Second, we sought a tissue matched line by comparing based on copy number signatures. In this instance, we performed clustering of the OVKATE line with all ovarian cancer cell lines based on copy number signatures using cosine similarity. OVKATE clustered with two other ovarian cell lines KUROMACHI and OVISE. These lines both showed highly correlated gene expression (r=0.79, p=0.02; and r=0.77, p=0.11) suggesting a relevant match in terms of chromosomal instability patterns and resulting gene expression changes.

## Discussion

Here we present CNpare, a tool for comparing cell line copy number profiles for the purpose of selecting optimal cell line models. CNpare is the first stand-alone tool to facilitate comparison of cancer cell lines based on high-resolution genome-wide copy number. This complements existing approaches based on low-resolution copy number, gene expression and methylation^3–13^. In addition, CNpare offers the option of comparing copy number profiles using copy-number signatures.

Despite showing excellent performance during benchmarking, this tool has two key limitations: 1) resolution - comparison is made at 500kb resolution (default bin size). While it is possible to increase the resolution up to 30kb, resolution beyond this is limited as the SNP6 technology underpinning the database does not facilitate higher resolution. 2) total absolute copy number as input. Total absolute copy number is currently required input and performance is dependent on the accuracy of the method used to compute it. We recommend using ASCAT^17^ as it matches the method used across the database.

Beyond matching cell line copy number profiles, CNpare can also be applied to other settings including: quality control - ensuring the sequenced copy number profile of a cell line matches the reference profile; assessing differences between cell line cultures - by estimating the percentage genome difference; and finding the best cell line model for a tumour profile - based on copy number profiles or copy number signatures. Our results demonstrate that CNpare can be used to select appropriately matched cancer cell line models, providing a valuable tool for improving cancer research in the context of studying chromosomal instability.

## Methods

### Data acquisition and curation

Precomputed absolute total copy number profiles for cell lines appearing in the Cancer Cell Line Encyclopaedia (CCLE)^18^ project and the Genomics of Drug Sensitivity in cancer (GDSC) project^19^ were downloaded from https://github.com/VanLoo-lab/ASCAT.sc^17^. Only cell lines with sufficient evidence of chromosomal instability were included in the CNpare database. To compute this, we smoothed normal segments by collapsing and merging near diploid segments, and then removed samples with less than 20 non-diploid aberrations. This reduced the number of available cell line profiles to 1,170.

### Data preprocessing

To enable comparison between two copy-number profiles, segmented profiles were converted into evenly sized bins across the genome (default 500 kb). To adjust for noise at copy number boundaries, a window-based smoothing procedure was used to align boundaries across samples. CNpare also provides the option to match samples based on the exposure to different types of chromosomal instability. To do this, we computed signature activities for the 7 ovarian signatures from copy-number profiles by applying the computational approach outlined in *Macintyre et al*.^*16*^. For running this approach, copy number data was initially formatted into a list of segment tables (one element of the list per sample). Then, values of six different features from each copy-number profile were computed, and then the linear combination decomposition function from YAPSA^20^ was used for identifying the signature activities in each sample.

### Comparison of copy-number profiles

Similarity between bin-level copy-number profiles was quantified using four different metrics: Pearson correlation coefficient, Manhattan distance, Euclidean distance and Cosine Similarity. The Pearson correlation test was computed using the *cor()* function from the *stats* R package (version 3.6.2), while the other metrics were directly calculated. As copy number signatures are compositional, only cosine similarity was used.

### Calculating percentage genome difference

Percentage genome difference was calculated at a copy number segment level and represents the fraction of the total human genome length where the segment value between the input profile and the matched profile is different. To compute this, copy number values of the input profile were recorded for each segment present in the matched cell line copy-number profile. If more than one segment of the input profile fell within a reference segment, these segments were merged by computing the median. Copy numbers were rounded to the nearest integer for computing differences. The total length of different segments was divided by the whole genome size, and the percentage reported.

### Comparison using copy-number signatures

Comparison based on copy number signatures was performed via clustering. In this case, CNpare reports similar cell lines based on shared cluster membership. k-means clustering was performed using the *akmeans()* function from the *akmeans* R package (version 1.1). This method automatically identifies the optimal number of clusters. Cosine similarity between the 7 copy number signatures was used as a distance metric.

### Comparing cell lines using other approaches

In order to benchmark CNpare against other approaches for choosing matched cell lines, we computed a number of variables representative of these alternative methods.

### Gene-level copy number

Gene positions were downloaded from UCSC Table Browser ^21^. We extracted the copy numbers in the transcription start of 27,212 protein-coding genes, and then compared gene-level copy numbers across cell lines using Pearson correlation.

### Chromosome arm copy number

We computed the whole arm chromosome copy number by averaging the values of all segments aligned to each chromosome arm. Then, we performed a Pearson correlation to compare CCLE and GDSC cell lines.

### Ploidy status

For each cell line, we computed the weighted mean copy number values of all segments to infer the overall ploidy. The relative weight of each segment depends on their length. Mean values were then rounded to the closest integer value, and the proportion of cell pairs matched by the ploidy status was then computed (81.57%, 248 cells).

### Gene-expression profiles

RNAseq FPKM gene expression data from 73 cell lines included in both CCLE and GDSC were downloaded from Cell Model Passports (https://cellmodelpassports.sanger.ac.uk/downloads). Gene expression levels were compared across cell lines using Pearson correlation using *cor*.*test*() from the stats R package (version 3.6.2).

### Testing suitability of OVKATE cell line matches

To validate the suitability of the cell lines matched with OVKATE we assessed correlation of gene expression and DNA methylation between the lines.

For gene expression, we downloaded log2 transformed TPM gene expression data for the protein coding genes from DepMap Public 21Q3 (https://depmap.org/portal/download/). We selected those genes which had an absolute log-fold change greater than 1 for OVKATE and used these genes to perform a Pearson correlation with PANC0203, KURAMOCHI and OVISE cell lines using *cor*.*test()* from the *stats* R package (version 3.6.2). Empirical p-values were calculated by dividing the number of cell lines with a higher r value than the observed r, by the total number of correlations made.

For DNA methylation, data from 1Kb upstream of gene transcription start sites (CCLE_RRBS_TSS1kb_20181022.txt.gz) was obtained from CCLE 2019^18^. Methylation profiles for cell lines EKVX, UO31, HOP62, MOLT3, OE21, JHUEM7, ML1, CL34, REC1 and OPM2 were removed due to insufficient data. Pearson correlation was calculated between all cell line methylation profiles using *cor*.*test()* from the *stats* R package (version 3.6.2). Empirical p-values were calculated by dividing the number of cell lines with a higher r value than the observed r, by the total number of correlations made.

## Availability and implementation

CNpare is available as an R package at https://github.com/macintyrelab/CNpare. All analysis performed in the manuscript can be reproduced via the code found at https://github.com/macintyrelab/CNpare_analyses

## Acknowledgements

We would like to thank Maxime Tarabichi and the Van Loo Lab for providing access to the ASCAT.sc processed CCLE and GDSC copy number profiles.

